# Role of the two flagellar stators in swimming motility of *Pseudomonas putida*

**DOI:** 10.1101/2022.08.03.502738

**Authors:** Veronika Pfeifer, Sönke Beier, Zahra Alirezaeizanjani, Carsten Beta

**Author notes:** Address correspondence to Carsten Beta,.

## Abstract

In the soil bacterium *Pseudomonas putida*, the motor torque for flagellar rotation is generated by the two stators MotAB and MotCD. Here, we construct mutant strains, in which one or both stators are knocked out and investigate their swimming motility in fluids of different viscosity and in heterogeneous structured environments (semisolid agar). Besides bright-field imaging of single cell trajectories and spreading cultures, dual color fluorescence microscopy allows us to quantify the role of the stators in forming *P. putida’s* three different swimming modes, where the flagellar bundle pushes, pulls, or wraps around the cell body. The MotAB stator is essential for swimming motility in liquids, while spreading in semisolid agar is not affected. Moreover, if the MotAB stator is knocked out, wrapped mode formation under low viscosity conditions is strongly impaired and only partly restored for increased viscosity and in semisolid agar. In contrast, when the MotCD stator is missing, cells are indistinguishable from the wild-type in fluid experiments, but spread much slower in semisolid agar. Analysis of the microscopic trajectories reveals that the MotCD knockout strain forms sessile clusters thereby reducing the number of motile cells, while the swimming speed is unaffected. Together, both stators ensure a robust wild-type that swims efficiently under different environmental conditions.

**IMPORTANCE:** Because of its heterogeneous habitat, the soil bacterium *Pseudomonas putida* needs to swim efficiently under very different environmental conditions. In this paper, we knocked out the stators MotAB and MotCD to investigate their impact on swimming motility of *P. putida*. While the MotAB stator is crucial for swimming in fluids, in semisolid agar both stators are sufficient to sustain a fast swimming phenotype and increased frequencies of the wrapped mode, which is know to be beneficial for escaping mechanical traps. However, in contrast to the MotAB knock-out, a culture of MotCD knock-out cells spreads much slower in the agar as it forms non-motile clusters that reduce the amount of motile cells.

---

**F**lagella-mediated swimming is one of the most common strategies of locomotion in the bacterial world. Powered by membrane embedded molecular motors, bacteria rotate their flagella to propel themselves and have evolved a wide variety of swimming strategies depending on the arrangement of flagella across the cell body (1). The flagellar movement is driven by stator complexes that rely on ion gradients across the membrane to generate the torque required for rotation. This machinery is dynamic with stators being constantly recruited and released from the motor, exchanging with a large pool of freely diffusing stators in the membrane, so that bacteria can flexibly adapt to changing environmental conditions (2). The widely studied model organism *Escherichia coli* operates a single type of stator, the proton-driven MotAB complex. It consist of a MotB dimer surrounded by five MotA proteins (3, 4). Depending on the mechanical load on the flagellum, up to eleven motAB stators can bind to the motor of *E. coli* (5, 6). However, numerous bacteria have more than one type of stator allowing them to fine-tune their motor function by, for example, changing the stator composition and recruiting the more favorable stator for a given task. *Shewanella oneidensis* has sodium-driven PomAB stators and proton-driven MotAB stators to adjust its motor function in the presence of different sodium concentrations (7). In *Vibrio parahaemolyticus*, two flagellar systems with two different types of stators have evolved. The polar flagellum is propelled by sodium-driven stators and the lateral flagella by proton-driven ones (8). In *Pseudomonas aeruginosa* the two stators MotAB and MotCD use proton motive force to power the rotation of a single flagellum. Both stators can promote swimming motility in aqueous environments, but only MotCD can maintain swimming under high viscosity conditions and swarming across surfaces (9, 10).

The closely related soil bacterium *Pseudomonas putida* also possesses the two stators MotAB and MotCD (11). However, *P. putida* does not swarm in a flagella-dependent manner (12) and is lophotrichously flagellated, having multiple flagella at one cell pole (13). *P. putida* displays a complex swimming pattern consisting of straight runs with two alternating speeds that are interrupted by stops and directional reversals (14). Three different run modes can be distinguished. When the flagellar bundle rotates counterclockwise (CCW), it pushes the cell body forward. When it rotates clockwise (CW) it can either pull the cell body or wrap around it, resulting in a slower swimming speed (15). Moreover, in the wrapped mode *P. putida* responds to chemoattractant gradients, which is considered to be a beneficial strategy to navigate its crowded natural habitat (16). The wrapped mode is known to also form in other polarly flagellated species (17), such as monopolarly flagellated *Shewanella putrefaciens* (18) and lophotrichously flagellated *Burkholderia* sp. RPE64 (19). More recently, it was also discovered in amphitrichous *Campylobacter jejuni* (20) and monotrichous *P. aeruginosa* (21).

For *P. putida* the question arises, if there is a specific role of the two different stators MotAB and MotCD for swimming motility and how the stators influence the swimming modes, especially the formation of the wrapped mode. It is assumed that the wrapped mode is formed due to changes in motor torque (15, 18). Recently, numerical simulations supported this conjecture, showing increased wrapped mode formation under higher torque (22). In this study, we knocked out the two stators MotAB and MotCD and analysed swimming motility under different environmental conditions to address these questions.

## RESULTS

### Construction of stator deletion mutants

*P. putida* has a large gene cluster for flagellar motility comprising the genes for the stator MotCD (PP4336-PP4335). Additionally the genome encodes a second copy of a stator outside this gene cluster, the MotAB (PP4905-PP4904) (11). Protein sequence alignments show a high similarity to the MotAB and MotCD stators in *P. aeruginosa* PAO1 (Figure S1). The MotA shares 81% identity, MotB 70% identity, MotC 83% identity and MotD 70% identity. We targeted these genes in *P. putida* to generate single and double deletion mutants. As a control, we rescued the mutants by reintroducing the genes. Swimming in aqueous environments and swimming agar assays confirmed a fully restored wild-type phenotype (Figure S2). For none of the mutants the growth rate was influenced, so that all cultures grow at a similar speed.

### MotAB is essential for swimming and wrapped mode formation under low viscosity conditions

To characterize the role of the two stators, we compared the swimming motility of the wild-type, the single mutants Δ*motAB* and Δ*motCD* and the double mutant Δ*motAB* Δ*motCD*. A visual inspection of swimming motility from shaking cultures with phase contrast microscopy revealed clear differences between the four strains. The double mutant Δ*motAB* Δ*motCD* is non-motile. Swimming motility of the Δ*motCD* mutant is indistinguishable from the swimming motility of wild-type cells, whereas the Δ*motAB* mutant shows strong deficiencies in swimming. Figure 1A shows a selection of the longest trajectories resulting from cell segmentation and tracking with a custom-made MATLAB software. While for the wild-type and the Δ*motCD* strain the trajectories mainly consist of long, straight runs that are interrupted by occasional abrupt turning events, the Δ*motAB* strain exhibits only short, erratic trajectories. When calculating the mean speed over the complete set of trajectories, we obtained 32.1 *µ*m/s ± 0.3 *µ*m/s for the wild-type, 6.6 *µ*m/s ± 0.2 *µ*m/s for the Δ*motAB* mutant, and 32.5 *µ*m/s ± 0.3 *µ*m/s for the Δ*motCD* mutant. Thus, the Δ*motAB* mutant swims more than four times slower than the wild-type, which is also reflected in a shift of the mean square displacement (MSD) towards lower values of the diffusivity, see Figure 1B.

**FIG 1.**
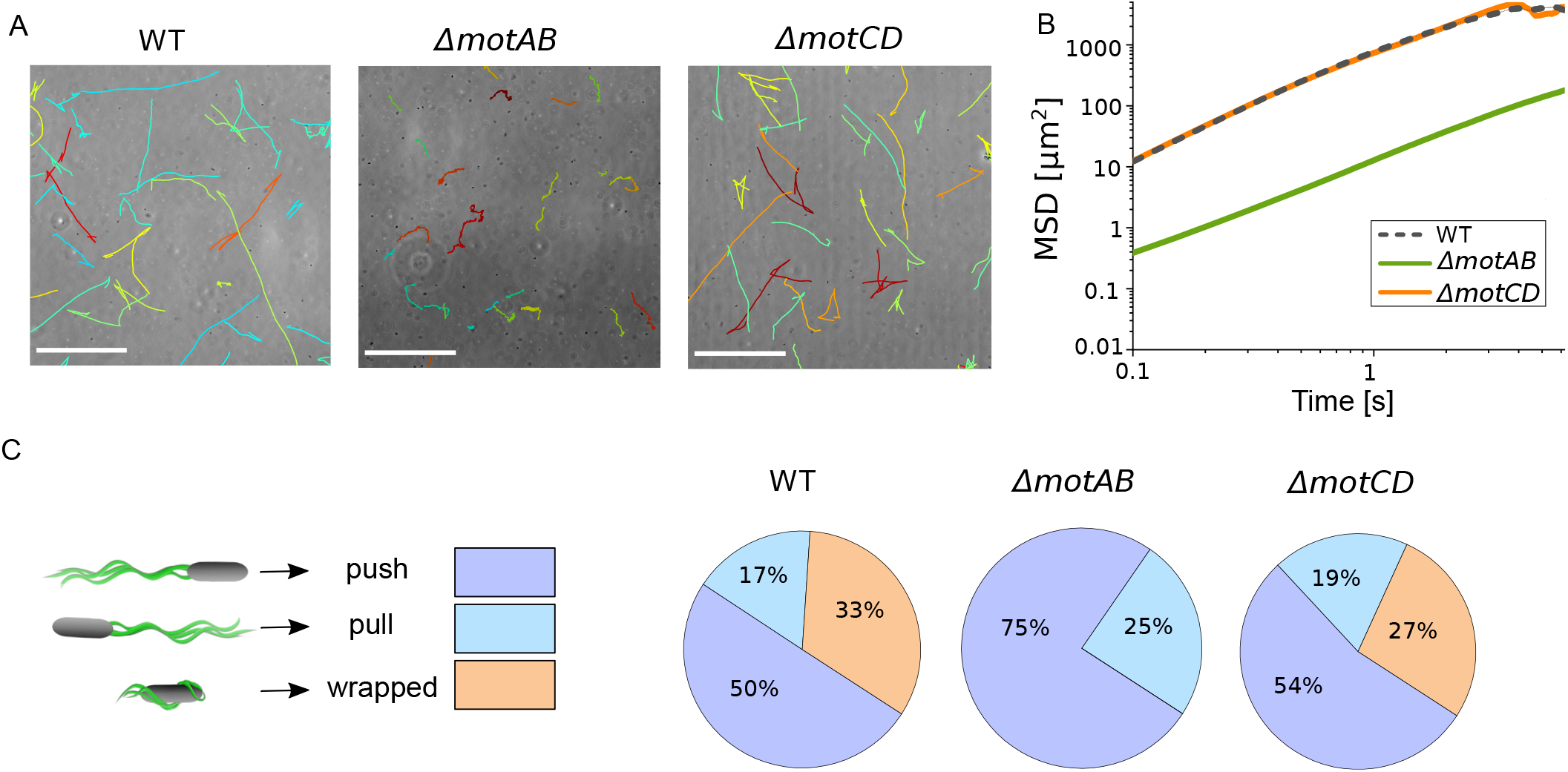
Swimming in aqueous environment reveals strong deficiencies of the Δ*motAB* mutant, whereas the Δ*motCD* mutant shows similar behavior as the wild-type strain. (A) Swimming trajectories for the wild-type and the Δ*motAB* and Δ*motCD* mutant strains. Scale bar is 100 *µ*m. (B) Mean square displacement over time. For the wild-type and the Δ*motCD* strain more than thousand trajectories were used, while 280 trajectories of the Δ*motAB* strain were included. The smaller sample size for the Δ*motAB* mutant is due to the decreased motility. (C) Pie charts showing the ratios of the different swimming modes contributing to swimming motility. We manually analyzed 441 runs for the wild-type strain, 216 runs for the Δ*motAB* mutant, and 268 runs for the Δ*motCD* mutant. The cartoons on the left-hand side show the three swimming modes push, pull and wrapped, the arrows indicate the swimming direction.

In order to distinguish between the three different run modes — push, pull, and wrapped — that are part of *P. putida*’s swimming strategy, we stained the flagella and visualized them by fluorescence microscopy with high spatial and temporal resolution. Figure 1C shows the frequency of the three run modes as they were observed during swimming of the wild-type, the Δ*motAB*, and the Δ*motCD* mutant strains. In agreement with the morphology of the trajectories observed by phase contrast imaging, the distributions of run modes for the wild-type and the Δ*motCD* mutant strain are similar, while the distribution for the Δ*motAB* mutant differs. Not only are the Δ*motAB* cells less motile in an aqueous environment, but they are also not able to form the wrapped mode. Surprisingly, the ratio of CW (pull and wrap mode) to CCW (push mode) rotation of the flagella is influenced as well: the Δ*motAB* mutant shows an increased amount of runs with CCW flagellar rotation.

### Increasing viscosity partly restores wrapped mode formation in the absence of MotAB

For other wrapped-mode forming bacteria it was shown that a more viscous surrounding medium leads to an increased portion of runs in the wrapped mode (18, 20). To test whether *P. putida* shows a similar behavior, we added Ficoll to swimming wild-type and stator mutant strains, to increase the viscosity and, thus, the load on the flagella. In particular, it was our aim to investigate whether the MotAB stator is strictly indispensable for wrapped mode formation, or whether there are conditions where also the Δ*motAB* knock-out mutant is able to form the wrapped mode.

In general, an increased Ficoll concentration led to decreased swimming speeds for all three strains (table 1). The decrease in swimming speed for the Δ*motCD* mutant was similar to the decrease in the wild-type strain. In contrast, the swimming speed of Δ*motAB* cells was already much lower in the absence of Ficoll. Adding Ficoll slowed the cells down even further, so that for Ficoll concentrations of 15% and higher, it was no longer possible to extract any trajectories. Nevertheless, the swimming modes could be still identified by fluorescence microscopy. We determined the swimming motility and flagellar configuration after adding 10%, 15%, and 20% of Ficoll. Under conditions of increased viscosity, the Δ*motAB* mutant showed a CW-CCW-ratio comparable to the wild-type and the Δ*motCD* mutant: for the Ficoll experiments the CW-CCW-ratio stays roughly constant for all strains, with the CCW push mode occurring between 50% and 65%, see Table S1. Figure 2 shows the distribution of the CW swimming modes, pull and wrap, at different Ficoll concentrations. For the wild-type and the Δ*motCD* strain, the portion of wrapped runs increases, until at 20% Ficoll, almost all CW runs are taking place in wrapped mode. Interestingly, also the Δ*motAB* mutant is able to form the wrapped mode under high viscosity conditions. However, at 20% Ficoll the amount of wrapped runs is decreasing again, in contrast to the wild-type and the Δ*motCD* cells.

**TABLE 1.**
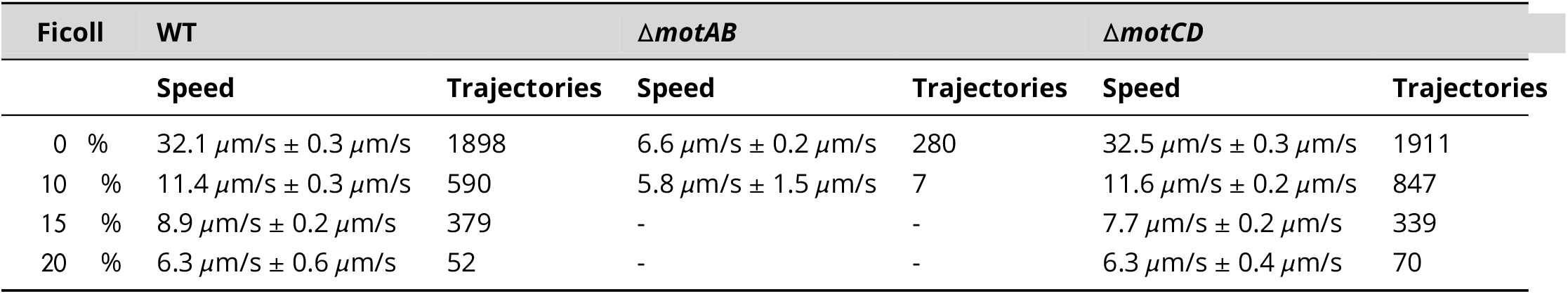
Mean speed calculated over the complete trajectories in different Ficoll concentrations. For the Δ*motAB* mutant motility is too low to extract the swimming speed at Ficoll concentrations of 15% or higher.

**FIG 2.**
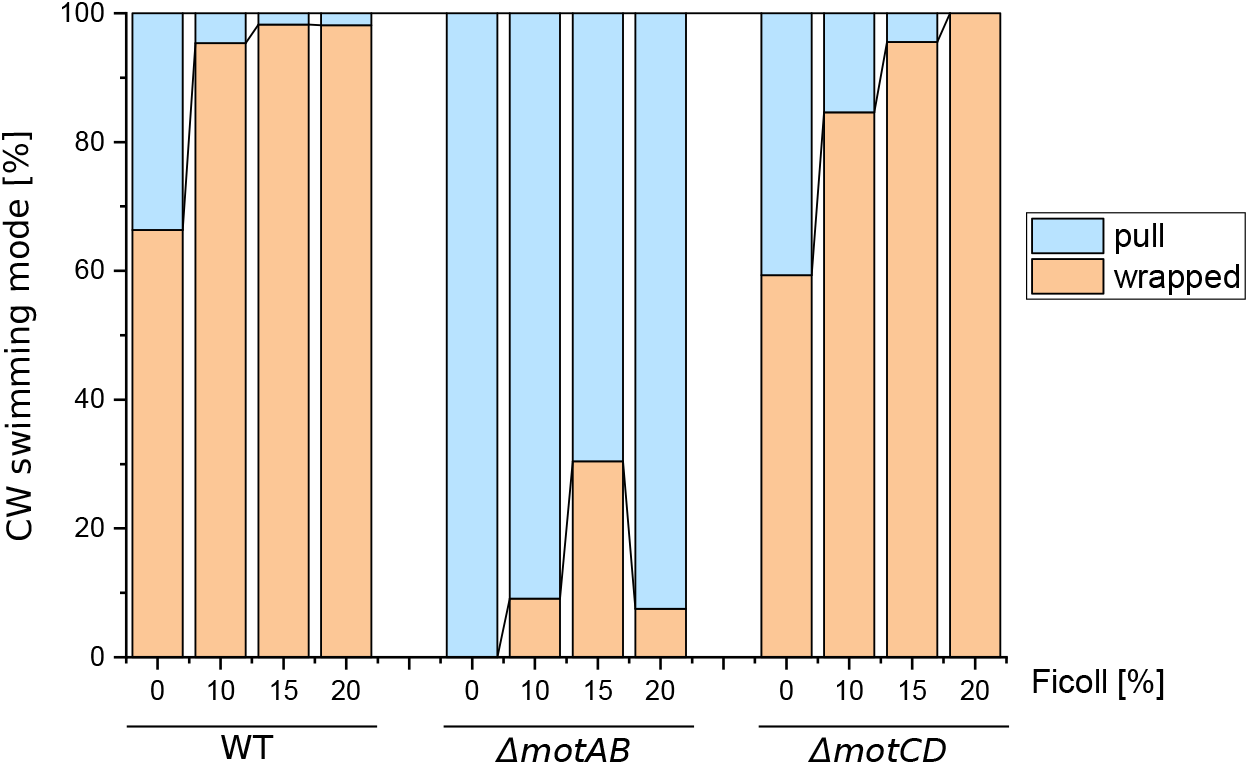
Influence of increased viscosity on the ratio of CW swimming modes pull and wrapped. In general, the stepwise increase in viscosity led to increased wrapped mode formation, even Δ*motAB* was able to form the wrapped mode under higher viscosity conditions. However, for the Δ*motAB* mutant the amount of wrapped mode increased only for up to 15% Ficoll and decreased again for 20% Ficoll. The ratio of the three swimming modes push, pull, and wrapped and the sample sizes of all experiments are listed in Table S1.

### In semisolid agar, Δ*motAB* mutant spreads similar to wild-type and outperforms Δ*motCD* mutant

In their natural soil habitat, *P. putida* cells have to navigate through strongly confined, narrow spaces of complex, irregular geometry. To elucidate the role of the two stators with respect to swimming motility in such complex environments, we have investigated the swimming of *P. putida* in semisolid agar. Semisolid agar is a randomly structured, close-meshed, porous network filled with fluid (23). We injected cells into 0.25% and 0.30% semisolid agar and quantified the macroscopic spreading of the growing culture over time, see Figure 3A. As expected, cells spread faster in 0.25% than in 0.30% agar due to the difference in pore size, see Figure 3B. For both agar concentrations the Δ*motAB* culture spreads almost as fast as the wild-type, whereas the Δ*motCD* culture spreads considerably slower. This cannot be explained based on the observed swimming behaviour in open fluids, where, in contrast to the spreading in agar, Δ*motAB* cells are barely motile and Δ*motCD* cells swims with similar speeds as the wild-type.

**FIG 3.**
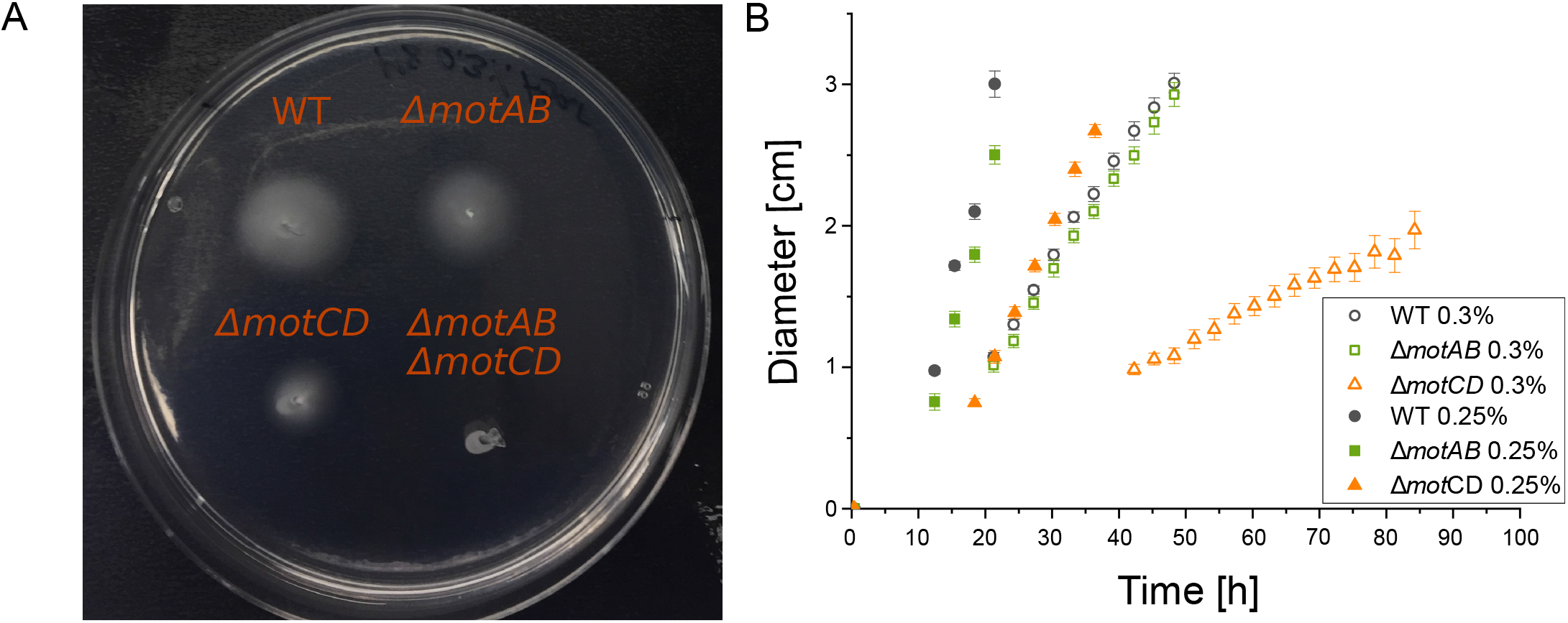
Spreading of bacterial cultures in swimming agar. (A) Spreading of the wild-type, the single mutants Δ*motAB* and Δ*motCD*, and the double mutant Δ*motAB* Δ*motCD* in a 0.30% semisolid agar plate. The double mutant is non-motile. (B) Time evolution of the diameter of wild-type (black), Δ*motAB* mutant (green), and Δ*motCD* mutant (orange) cultures in 0.25% (filled symbols) and 0.30% (open symbols) semisolid agar.

### Wild-type and both stator mutants show similar speed distributions in agar but wrapped mode formation is less frequent in Δ*motAB* mutant

To understand the different spreading efficiencies of wild-type and mutant strains in semisolid agar, we imaged and analyzed the motility of single cells in the agar matrix. Due to the narrow fibrous meshwork, swimming in semisolid agar differs from swimming in an open aqueous environment, see Movie S1 for an example. In particular, the length of straight runs is restricted to the free path length set by the geometry of the porous medium, resulting in more frequent interruptions of the runs as cells collide with or get stuck in the meshwork and have to reorient to escape. Figure 4A shows a selection of the longest swimming trajectories that we extracted from phase contrast imaging experiments of cells swimming in semisolid agar. Using an adapted version of our MATLAB-based cell tracking software, we determined the swimming speeds of the runs for the three different strains in agar (Fig. 4B). Comparing the speed distributions of the three strains, it turns out that the differences between them are less pronounced in agar than in aqueous fluid, where the mean speed is much lower for the Δ*motAB* mutant than for the wild-type and the Δ*motCD* mutant strain (Table 1). In the agar, in contrast, the mean speeds are much closer, with the Δ*motCD* mutant swimming slightly faster and the Δ*motAB* mutant slightly slower than the wild-type. The mean speed per run is 23.85 *µ*m/s ± 0.05 *µ*m/s for the wild-type, 21.44 *µ*m/s ± 0.04 *µ*m/s for the Δ*motAB* mutant, and 27.18 *µ*m/s ± 0.12 *µ*m/s for the Δ*motCD* mutant (please note that here the speed values are calculated for the run episodes only, while in Table 1 averages are taken over entire trajectories). Based on this observation, we cannot explain the slower spreading of the Δ*motCD* mutant as compared to the Δ*motAB* and wild-type cultures in agar. However, it indicates an up-regulation of the MotCD stator in agar, leading to a Δ*motAB* mutant that swims almost as fast as the wild-type and the Δ*motCD* mutant.

**FIG 4.**
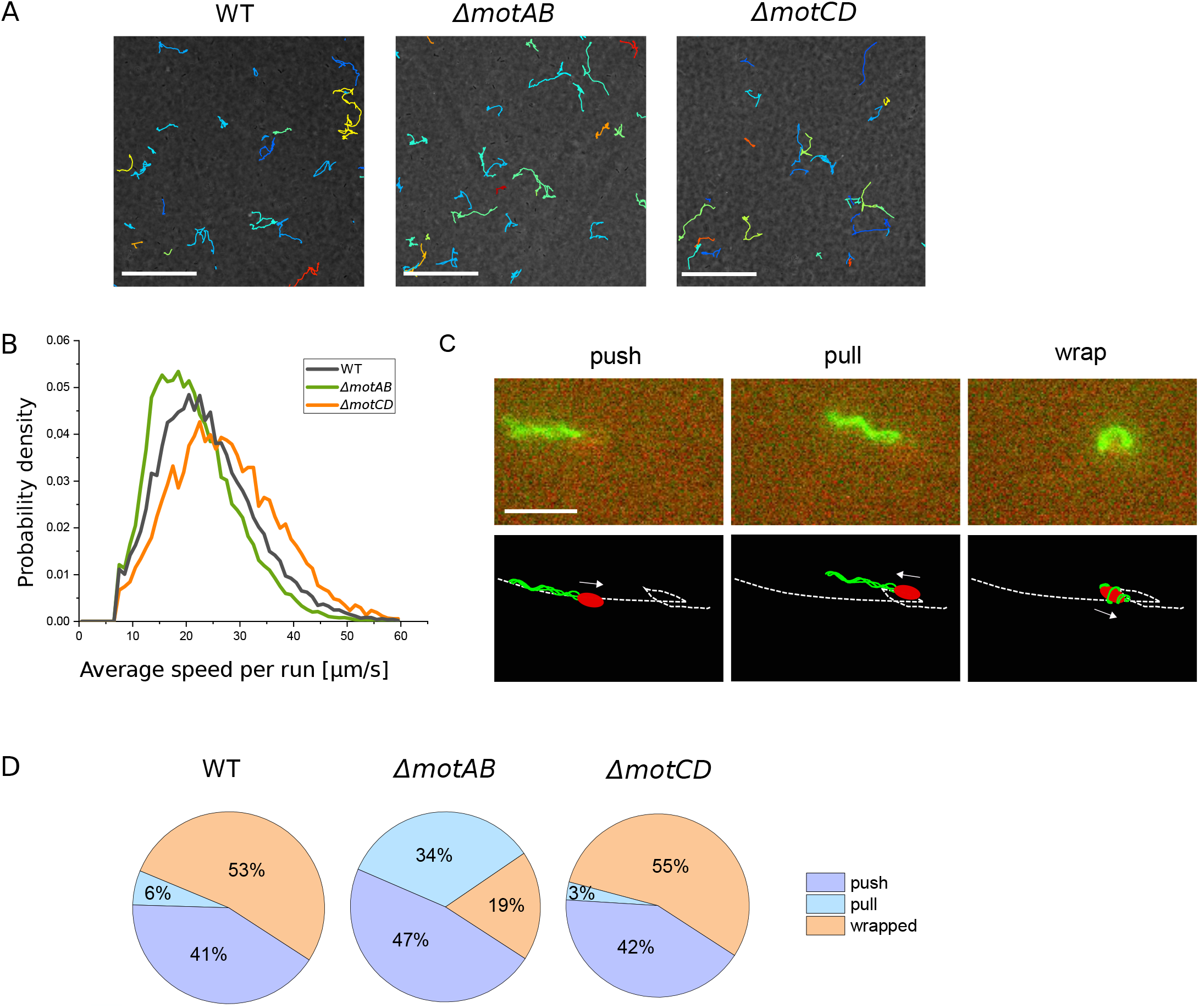
Single cell motility in 0.25% swimming agar. (A) Swimming trajectories in semisolid agar. Scale bar is 100 *µ*m. (B) Distribution of mean run speeds in semisolid agar. (C) The three swimming modes in semisolid agar shown for luorescently labeled wild-type cells, see also the corresponding supplementary Movie S2. The arrows indicate the swimming direction. Scale bar is 5 *µ*m. (D) Pie charts showing the frequency of swimming modes. We manually analyzed 327 runs for the wild-type, 241 runs for the Δ*motAB* mutant, and 189 runs for the Δ*motCD* mutant.

To determine the frequencies of the push, pull, and wrapped modes during swimming in semisolid agar, we labeled the flagella and the cell body with two different fluorescent dyes. Using dual color fluorescence microscopy, we simultaneously imaged their position and relative orientation to identify the swimming mode of each run for wild-type, Δ*motAB*, and Δ*motCD* mutant strains swimming in semisolid agar (Fig. 4C and Movie S2-S4). The staining experiments revealed an increased wrapped mode formation for all three strains in comparison to swimming in aqueous fluid (Fig. 4D). Surprisingly, also in semisolid agar, the distribution of swimming modes for the wild-type and the Δ*motCD* mutant are similar, while a different distribution with a smaller portion of the wrapped mode is observed for the Δ*motAB* mutant. This is consistent with the observations in viscous fluids but does not explain the different macroscopic spreading dynamics of the Δ*motCD* culture as compared to the wild-type and Δ*motAB* cultures in semisolid agar (Fig. 3A).

### Cluster formations of Δ*motCD* mutant in semisolid agar results in fewer motile cells, thereby decreasing spreading of the culture

In contrast to the wild-type and the Δ*motAB* cultures, the number of motile cells of the Δ*motCD* mutant strain decreased over time in the swimming agar assay. Instead, we noticed the formation of growing clusters of immobile cells. To analyze the cluster formation in more detail, we recorded the Δ*motCD* culture for several hours and compared it to similar imaging experiments with wild-type and Δ*motAB* cultures in semisolid agar. These recordings show that motile cells occasionally stop before they divide, most likely because they get trapped. For wild-type and Δ*motAB* mutant cells at least one, but oftentimes both daughter cells move on after division (Fig. 5A). In contrast, the Δ*motCD* mutant cells mostly do not become motile again after division, so that both daughter cells remain immobile. These cells are still alive and by further divisions of the daughter cells the above-mentioned cell clusters are formed (Fig. 5B). Cluster formation of the Δ*motCD* mutant cells was more pronounced in 0.3% agar. Here, the daughter cells remained immobile after most of the cell divisions we observed. In 0.25% agar, in addition to the cluster forming division events, also cell divisions resulting in motile daughter cells were seen. Thus, the number of motile Δ*motCD* cells was higher in 0.25% than in 0.3% agar, but still lower than in the cultures of wild-type and Δ*motAB* mutant strains, resulting in a strongly reduced spreading of the Δ*motCD* culture (Fig. 3).

**FIG 5.**
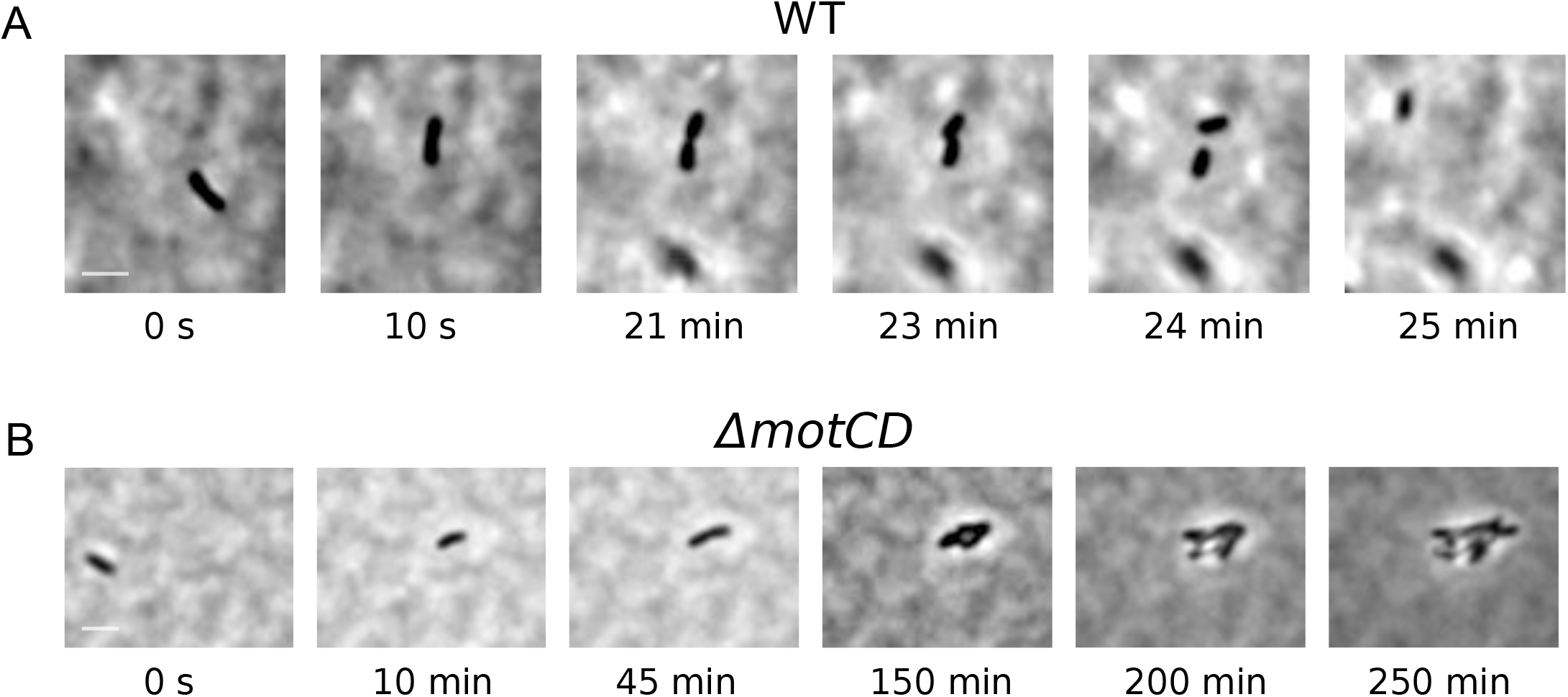
Cell division of trapped cells in semisolid agar. (A) Wild-type cells move on after division. (B) Δ*motCD* mutant cells remain sessile and form growing clusters. See also the corresponding supplementary Movies S5 and S6. Scale bar is 5 *µ*m.

## DISCUSSION

We examined swimming motility of wild-type and stator mutant cells of the soil bacterium *P. putida* in different environments. In homogeneous fluids, the MotAB stator is crucial to promote swimming motility, so that the movement of Δ*motAB* mutant cells in aqueous fluids is strongly impaired. In contrast, in a heterogeneous, structured agar environment, both stators can promote swimming motility but the MotCD stator, in addition, prevents cells from forming sessile clusters. Taken together, the two stators ensure robust swimming motility of *P. putida* under different environmental conditions.

Along with the strong swimming deficiencies in Δ*motAB* mutant cells, also the ratio of CCW to CW episodes of flagellar rotation is altered. While the wild-type and the Δ*motCD* mutant show approximately equal ratios of CCW (push) and CW (pull and wrapped) swimming modes, only 25% of the Δ*motAB* mutant cells exhibited CW flagellar rotation (only pull, no wrapped mode) when swimming under low viscosity conditions (Fig. 1C). For *E. coli* it is known that CW rotating motors generate a lower torque at intermediate load than CCW rotating motors (24), which may be related to an asymmetry in the load dependent stator binding between CW and CCW rotating motors (6). If the same is true for *P. putida*, the increased portion of CCW runs may indicate that, under conditions where swimming motility is strongly impaired, cells adapt their locomotion strategy such that they preferentially rely on CCW rotation of their flagella to generate sufficient torque for propagation. Alternatively, the MotAB knockout might directly influence the chemotaxis pathway leading to an altered CW bias. This is supported by observation in *P. aeruginosa* showing that MotC interacts with the diguanylate cyclase SadC, which may lead to an increased production of the key motility regulator cyclic diguanylate (25).

For the Δ*motAB* mutant not only the relative frequencies of CCW and CW swimming modes are altered but also the ratio of pull and wrapped modes, that may both occur during CW rotation of the flagellar motors, is drastically changed. For wild-type and Δ*motCD* mutant cells both pull and wrapped mode runs are frequently observed (Fig. 1C), with an increasing portion of wrapped runs for higher viscosities (Fig. 2), similar to previous observations in *S. putrefaciens* (18). In contrast, under low viscosity conditions, the Δ*motAB* mutant does not exhibit any wrapped mode formation at all (Fig. 1C). Only for increased viscosity and in an agar meshwork, a low fraction of wrapped runs is observed for this mutant strain (Figs. 2 and 4C). Thus, wrapped mode formation is not restricted to one type of stator. Also the MotCD stator is sufficient to form the wrapped mode under increased load, even though less efficiently. It was previously conjectured that wrapped mode formation results from an instability of the pulling bundle that is triggered by an increase in motor torque (18, 15). Numerical simulations of lophotrichously flagellated bacteria have confirmed this torque dependent filament wrapping (22). We thus assume that, under low viscosity conditions, Δ*motAB* mutant cells are not able to form the wrapped mode because not enough torque is generated by the MotCD stators that are less active in fluids, cf. also the low swimming speeds of the Δ*motAB* mutant cells. Under increased load, presumably more MotCD stators are recruited to the motor, increasing the torque to such an extent that occasional wrapped mode formation can be observed. This agrees with the observation that also in *P. aeruginosa*, less MotCD stators bind to the motor during operation in a low viscosity environment (26).

Our findings for *P. putida* differ from earlier observations reported for Δ*motAB* and Δ*motCD* mutants in *P. aeruginosa*. For swimming in fluids the speed of the Δ*motAB* knock-out in *P. aeruginosa* is only slightly reduced compared to the wild-type (9), in contrast to the strong effect we observed for the *P. putida* Δ*motAB* mutant (Table 1). Furthermore, for *P. aeruginosa* it was shown that under increased load on the flagella, more MotCD units are recruited to the motor (26). An increased viscosity thus only leads to a decrease in motilty for the Δ*motAB* mutant and the wild-type while it abolishes motility in the Δ*motCD* mutant for 15% Ficoll (10). Therefore, MotCD is assumed to be required for generating higher torque in *P. aeruginosa*. In contrast, for *P. putida* we observe that, under high viscosity conditions, the Δ*motAB* mutant exhibits only little wrapped mode formation and its swimming speed remains low, while Δ*motCD* cells are comparable to the wild-type in terms of wrapped mode formation and swimming speeds (Fig. 2 and Table 1).

Only when changing the environmental conditions to a heterogeneous agar meshwork, we see that the Δ*motAB* mutant can outperform the Δ*motCD* mutant by spreading faster in semisolid agar (Fig. 3). This agrees with earlier observations in *P. aeruginosa* (9). However, the slower spreading of the *P. putida* Δ*motCD* culture is not caused by a decreased swimming speed. Surprisingly, the Δ*motCD* mutant swims at similar speeds as the Δ*motAB* and wild-type strain (Fig. 4C) but less motile cells are observed due to the formation of sessile clusters (Fig. 5). The mechanism behind the cluster formation is yet unknown. It may be related to the surface sensing properties of MotAB, which were observed for *P. aeruginosa* (27). We assume that cluster formation is the main reason for slower spreading of the Δ*motCD* culture. Effectively, clustering decreases the amount of newly formed motile daughter cells, which will influence the cell culture spreading (28). Specifically, we assume that the observed linear increase in culture radius (Fig. 3) results from an interplay of reproduction and motility, as was previously proposed (23), where a reduced growth rate of motile cells due to clustering will lead to a decreased slope in the linear time profile of the culture radius.

The natural habitat of *P. putida* is a heterogeneous, crowded soil environment. While earlier experiments in larger confined spaces, such as between parallel plates or cylindrical obstacles, revealed only slight modulations of the swimming pattern known from uniform aqueous liquid (29, 30), our present experiments in agar can be seen as a first step to mimic a more complex dense environment. The increased wrapped mode formation in agar that we observed for wild-type as well as for both stator mutant strains agrees with earlier observations that the wrapped mode typically occurs under external confinement and most likely serves as a strategy of flagellated cells to escape from mechanical traps (18, 19) or to enhance environmental spreading (31). It also fits the prediction that *P. putida* preferentially uses this slow swimming mode to navigate its natural environment (16, 22). How bacteria sense mechanical confinement and trapping remains an open question that will be addressed in subsequent studies. Furthermore, we aim to investigate in future experiments how the pore size and other geometric parameters of the environment influence the spreading dynamics in inhomogenous environments. For this purpose not only agar substrates will be used but also custom-made microfluidic chambers and other soft materials, such as hydrogels, that have already been successfully applied to describe the motility of *E. coli* in a complex, confined matrix (32, 33).

## MATERIALS AND METHODS

### Cell culture

We used the strain *P. putida* KT2440 FliC_*S*267*C*_. In a previous study, the flagellin protein FliC was genetically modified by exchanging the serine 267 with a cysteine in order to fluorescently label the flagella. It was shown that this exchange does not influence motility (15). We refer to this strain as wild-type. *P. putida* was grown in shaking culture at 30^°^C and the required *E. coli* strains for cloning at 37^°^C. For cloning and the swimming agar assay cells were inoculated in lysogeny broth. For flagellar staining experiments cells were grown over night in tryptone broth (10 g/l tryptone, Applichem, 5 g/l NaCl).

### Construction of vectors and strains

All required oligonucleotides and plasmids used in this study are listed in Table S2. To knock-out the stator genes *motAB* or *motCD*, for each stator its up- and downstream region were amplified and cloned into the suicide vector pNPTS138-R6K via Gibson assembly and transformed into *E. coli* DH5*αλ*pir competent cells. The recombinant vector was transformed into the donor cells *E. coli* WM3064 and further via conjugation to *P. putida*. In *P. putida*, we selected for sequential double homologous recombination (34). The reintroduction of the knocked-out genes for complementation was performed in the same way.

### Swimming experiments in luids

Fluid experiments were carried out in *µ*-Slide VI 0.1 channels (Ibidi). To increase viscosity, Ficoll 400 was added from 10% up to 20% (w/v).

### Staining experiments

Flagella were stained with the fluorescent dye Alexa 488 C5-maleimide (Thermo Fisher Scientific). The staining was carried out as published before in Hintsche et al. (15). For agar experiments we additionally stained the cell body with 10 *µ*l FM 4–64 (Thermo Fisher Scientific; 1 *µ*g/*µ*l in DMSO). It was added before the last washing step. Previous experiments showed that the double staining does not affect the swimming modes.

### Swimming agar assay

The swimming agar assay was carried out according to Ha et al. (35) with 0.3% and 0.25% semisolid agar. In short, M8 (42.4 g/l disodium phosphate dihydrate, 15 g/l monopotassium phosphate, 2.5 g/l sodium chloride) agar plates with 0.2% glucose, 0.5% casamino acids and 1 mM magnesium sulfate as final concentration were poured into petri dishes. Plates solidify over four hours. A small volume of cells was injected into the agar. Recordings of stained cells were carried out after additional four hours. After this time, the cells have adapted to the new environment and have spread away from the injection point where the agar is destroyed. We could not wait for more than four hours, because afterwards we barely find any stained cells as newly formed flagella which are built during cell division are not stained. Phase contrast recordings were performed on the next day. For the analysis of the culture spreading in semisolid agar standard petri dishes were used. For analysis of single cells FluoroDish Cell Culture Dishes with a glass bottom were used.

### Microscopy

Microscopy was carried out with an inverted microscope (Olympus IX71) with a blue LED as excitation source (4.8 W optical output power, 470 nm) for fluorescence microscopy and a white LED as light source for phase contrast. The ORCA-Fusion BT Digital CMOS camera from Hamamatsu was used for recordings together with the Hokawo software (Hamamatsu). Phase contrast was recorded 30 *µ*m over the bottom surface with a 20x UPLFLN-PH objective (Olympus) and 20 frames per seconds to analyze single cell swimming motility. To identify cluster formation of the Δ*motCD* mutant in the swimming agar assay, we recorded the culture for 15 hours with 3 frames per minutes and compared it to similar imaging experiments of wild-type and Δ*motAB* cultures. Fluorescence images were recorded close to the surface with 100 frames per second using a 60x UPLFLN-PH objective (Olympus). For the double fluorescence staining of the flagella and the cell body in the agar experiments the W-VIEW GEMINI Image Splitting Optics (Hamamatsu) was used. Fluorescence images were processed with ImageJ as described in Hintsche et al. (15). The frequency of swimming modes was determined by counting the number of observed modes.

### Processing with MATLAB

For segmentation and cell tracking the code described in Theves et al. (14) was used, which is based on Crocker and Grier (36). For measurements in bulk, the tested filter parameters also remain the same. The trajectories with an average velocity < 10 *µ*m/s, a displacement < 5 *µ*m, a length less than 2 s and the trajectories, which are highly curved, are therefore filtered out, see (14). For measurements in agar, on the other hand, only very short trajectories (< 1.5 s) and trajectories with very low displacement (< 1 *µ*m) are filtered out, since the velocities in agar are generally lower and bacteria do not move as much and as straight as in the bulk. To identify runs (Fig. 4) we use two conditions. All those phases of a track with a current speed that is below the threshold value of 7 *µ*m/s are not classified as runs. These phases may be called traps, following the wording of Bhattacharjee and Datta (32). As a second condition, time points of the tracks are excluded from run phases if they are located between two traps where the start point of the second trap is less than 2 *µ*m away from the end point of the previous trap. Two traps that are very close to each other thus merge into a single larger trap. This second condition had to be introduced because from visual inspection it became obvious that larger traps were frequently split into several smaller trap events when only the first condition was used. All time points that are not traps are classified as runs.

## SUPPLEMENTAL MATERIAL

**FIGURE S1**. Protein sequence alignment for the stators MotAB and MotCD in *P. putida* and *P. aeruginosa*. Alignment was made with BLAST (https://blast.ncbi.nlm.nih.gov/).

**FIGURE S2**. Complementation analysis for *motAB* and *motCD*. (A) Complementation of MotAB leads to a very motile phenotype in aqueous environments. The swimming motility of Δ*motAB* motAB cells is comparable with the wild-type swimming motility. (B) Complementation of MotCD in the Δ*motCD* mutant leads to increased spreading in agar comparable to the spreading of wild-type cells. Due to complementation of MotAB in the Δ*motAB* Δ*motCD* double mutant the strain shows a similar phenotype like the Δ*motCD* mutant.

**TABLE S1**. Frequency of swimming modes for different Ficoll concentrations. For each experiment the sample size is listed. For the Δ*motAB* mutant it is smaller due to the decreased motility. However, for 0% and 20% Ficoll we analysed a bigger amount of measurements to get a comparable sample size.

**TABLE S2**. Oligonucleotides and Plasmids used for molecular cloning.

**MOVIE S1**. Comparison of swimming in liquid and swimming in agar.

**MOVIE S2**. Dual color fluorescence imaging of wild-type swimmer in agar.

**MOVIE S3**. Dual color fluorescence imaging of Δ*motAB* mutant in agar.

**MOVIE S4**. Dual color fluorescence imaging of Δ*motCD* mutant in agar.

**MOVIE S5**. Cell division of trapped wild-type cell. One frame was made every 50 seconds.

**MOVIE S6**. Cell division of trapped Δ*motCD* mutant cell forming a cell cluster. One frame was made every 20 seconds.

## ACKNOWLEDGMENTS

We are grateful to Kai Thormann for providing the pNPTS138-R6KT plasmid and the *E. coli* strains for molecular cloning. Furthermore, we thank Sven Flemming and Marc Erhardt for helpful disussions. This research has been partially funded by the Deutsche Forschungsgemeinschaft (DFG) – Project-ID 318763901 – SFB1294 and – Project-ID 443369470 – BE 3978/13-1. We also acknowledge financial support by the Bundesministerium für Bildung und Forschung (BMBF) in the framework of “Ideenwettbewerb-Neue Produkte für die Bioökonomie” (Grant No. 031B0653A).

## REFERENCES

1. Grognot M, Taute KM. 2021. More than propellers: how flagella shape bacterial motility behaviors. Curr Opin Microbiol 61:73–81. doi:10.1016/j.mib.2021.02.005.

2. Leake MC, Chandler JH, Wadhams GH, Bai F, Berry RM, Armitage JP. 2006. Stoichiometry and turnover in single, functioning membrane protein complexes. Nature 443 (7109):355–358. doi: 10.1038/nature05135.

3. Deme JC, Johnson S, Vickery O, Aron A, Monkhouse H, Gri ths T, James RH, Berks BC, Coulton JW, Stansfeld PJ, Lea SM. 2020. Structures of the stator complex that drives rotation of the bacterial flagellum. Nat Microbiol 5 (12):1553–1564. doi:10.1038/s41564-020-0788-8.

4. Santiveri M, Roa-Eguiara A, Kühne C, Wadhwa N, Hu H, Berg HC, Erhardt M, Taylor NMI. 2020. Structure and Function of Stator Units of the Bacterial Flagellar Motor. Cell 183 (1):244–257.e16. doi: 10.1016/j.cell.2020.08.016.

5. Lele PP, Hosu BG, Berg HC. 2013. Dynamics of mechanosensing in the bacterial flagellar motor. Proc Natl Acad Sci 110 (29):11839–11844. doi:10.1073/pnas.1305885110.

6. Tipping MJ, Delalez NJ, Lim R, Berry RM, Armitage JP. 2013. Load-Dependent Assembly of the Bacterial Flagellar Motor. mBio 4 (4):1–6. doi:10.1128/mbio.00551-13.

7. Paulick A, Koerdt A, Lassak J, Huntley S, Wilms I, Narberhaus F, Thormann KM. feb 2009. Two different stator systems drive a single polar flagellum in Shewanella oneidensis MR-1. Mol Microbiol 71 (4):836–850. doi:10.1111/j.1365-2958.2008.06570.x.

8. Atsumi T, McCartert L, Imae Y. 1992. Polar and lateral flagellar motors of marine Vibrio are driven by different ion-motive forces. Nature 355:182–184. doi:10.1038/355182a0.

9. Doyle TB, Hawkins AC, McCarter LL. 2004. The complex flagellar torque generator of Pseudomonas aeruginosa. J Bacteriol 186 (19):6341–6350. doi:10.1128/JB.186.19.6341-6350.2004.

10. Toutain CM, Zegans ME, O’Toole GA. 2005. Evidence for two flagellar stators and their role in the motility of Pseudomonas aeruginosa. J Bacteriol 187 (2):771–777. doi:10.1128/JB.187.2.771-777.2005.

11. Martínez-García E, Nikel PI, Chavarría M, de Lorenzo V. 2014. The metabolic cost of flagellar motion in Pseudomonas putidaKT2440. Environ Microbiol 16 (1):291–303. doi:10.1111/1462-2920.12309.

12. Matilla MA, Ramos JL, Duque E, De Dios Alché J, Espinosa-Urgel M, Ramos-González MI. 2007. Temperature and pyoverdine-mediated iron acquisition control surface motility of Pseudomonas putida. Environ Microbiol 9 (7):1842–1850. doi:10.1111/j.1462-2920.2007.01286.x.

13. Harwood CS, Fosnaugh K, Dispensa M. 1989. Flagellation of Pseudomonas-Putida and Analysis of Its Motile Behavior. J Bacteriol 171 (7):4063–4066.

14. Theves M, Taktikos J, Zaburdaev V, Stark H, Beta C. 2013. A bacterial swimmer with two alternating speeds of propagation. Biophys J 105 (8):1915–1924. doi:10.1016/j.bpj.2013.08.047.

15. Hintsche M, Waljor V, Großmann R, Kühn MJ, Thormann KM, Peruani F, Beta C. 2017. A polar bundle of flagella can drive bacterial swimming by pushing, pulling, or coiling around the cell body. Sci Reports 7 (1):1–10. doi:10.1038/s41598-017-16428-9.

16. Alirezaeizanjani Z, Großmann R, Pfeifer V, Hintsche M, Beta C. 2020. Chemotaxis strategies of bacteria with multiple run-modes. Sci Adv 6 (22):1–7. doi:10.1126/sciadv.aaz6153.

17. Thormann KM, Beta C, Kühn MJ. 2022. Wrapped Up: The Motility of Polarly Flagellated Bacteria. Annu Rev Microbiol 76:349–67.

18. Kühn MJ, Schmidt FK, Eckhardt B, Thormann KM. 2017. Bacteria exploit a polymorphic instability of the flagellar filament to escape from traps. Proc Natl Acad Sci United States Am 114 (24):6340–6345. doi:10.1073/pnas.1701644114.

19. Kinosita Y, Kikuchi Y, Mikami N, Nakane D, Nishizaka T. 2018. Unforeseen swimming and gliding mode of an insect gut symbiont, Burkholderia sp. RPE64, with wrapping of the flagella around its cell body. ISME J 12 (3):838–848. doi:10.1038/s41396-017-0010-z.

20. Cohen EJ, Nakane D, Kabata Y, Hendrixson DR, Nishizaka T, Beeby M. 2020. Campylobacter jejuni motility integrates specialized cell shape, flagellar filament, and motor, to coordinate action of its opposed flagella. PLoS pathogens 16 (7):e1008620. doi: 10.1371/journal.ppat.1008620.

21. Tian M, Wu Z, Zhang R, Yuan J. 2022. A new mode of swimming in singly flagellated Pseudomonas aeruginosa. Proc Natl Acad Sci United States Am 119:1–10. doi:10.1073/pnas.2120508119/-/DCSupplemental.Published.

22. Park J, Kim Y, Lee W, Lim S. 2022. Modeling of lophotrichous bacteria reveals key factors for swimming reorientation. Sci Reports p 1–12. doi:10.1038/s41598-022-09823-4.

23. Licata NA, Mohari B, Fuqua C, Setayeshgar S. 2016. Diffusion of Bacterial Cells in Porous Media. Biophys J 110 (1):247–257. doi: 10.1016/j.bpj.2015.09.035.

24. Yuan J, Fahrner KA, Turner L, Berg HC. 2010. Asymmetry in the clockwise and counterclockwise rotation of the bacterial flagellar motor. Proc Natl Acad Sci 107 (29):12846–12849. doi: 10.1073/pnas.1007333107.

25. Baker AE, Webster SS, Diepold A, Kuchma SL, Bordeleau E, Armitage JP, O’Toole GA. 2019. Flagellar Stators Stimulate c-di-GMP Production by Pseudomonas aeruginosa. J Bacteriol 5 (33):4–5.

26. Wu Z, Tian M, Zhang R, Yuan J. 2021. Dynamics of the two stator systems in the flagellar motor of Pseudomonas aeruginosa studied by a bead assay. Appl Environ Microbiol 87 (23). doi:10.1128/AEM.01674-21.

27. Laventie BJ, Sangermani M, Estermann F, Manfredi P, Planes R, Hug I, Jaeger T, Meunier E, Broz P, Jenal U. 2019. A Surface-Induced Asymmetric Program Promotes Tissue Colonization by Pseudomonas aeruginosa. Cell Host Microbe 25 (1):140–152.e6. doi: 10.1016/j.chom.2018.11.008.

28. Wolfe AJ, Berg HC. 1989. Migration of bacteria in semisolid agar. Proc Natl Acad Sci United States Am 86 (18):6973–6977. doi: 10.1073/pnas.86.18.6973.

29. Theves M, Taktikos J, Zaburdaev V, Stark H, Beta C. 2015. Random walk patterns of a soil bacterium in open and confined environments. EPL 109 (2):28007. doi:10.1209/0295-5075/109/28007.

30. Raatz M, Hintsche M, Bahrs M, Theves M, Beta C. 2015. Swimming patterns of a polarly flagellated bacterium in environments of increasing complexity. Eur. Phys. J. Spec. Top. 224 (7):1185–1198. doi: 10.1140/epjst/e2015-02454-3.

31. Kühn MJ, Edelmann DB, Thormann KM. 2022. Polar flagellar wrapping and lateral flagella jointly contribute to Shewanella putrefaciens environmental spreading. Environ Microbiol p 1–13. doi: 10.1111/1462-2920.16107.

32. Bhattacharjee T, Datta SS. 2019. Bacterial hopping and trapping in porous media. Nat Commun 10 (1):2–10. doi:10.1038/s41467-019-10115-1.

33. Bhattacharjee T, Datta SS. 2019. Confinement and activity regulate bacterial motion in porous media. Soft Matter 15 (48):9920–9930. doi: 10.1039/c9sm01735f.

34. Kühn MJ, Schmidt FK, Farthing NE, Rossmann FM, Helm B, Wilson LG, Eckhardt B, Thormann KM. 2018. Spatial arrangement of several flagellins within bacterial flagella improves motility in different environments. Nat Commun 9 (1):1–12. doi:10.1038/s41467-018-07802-w.

35. Ha DG, Kuchma SL, O’Toole GA. 2014. Plate-Based Assay for Swimming Motility in Pseudomonas aeruginosa, vol 1149. Springer Science+Business Media. doi:10.1007/978-1-4939-0473-0.

36. Crocker J, Grier D. 1995. Methods of Digital Video Microscopy for Colloidal Studies | Elsevier Enhanced Reader. J Colloid Interface Sci 179 (179):298–310. https://doi.org/10.1006/jcis.1996.0217.

